# Individual variability in behavior and functional networks predicts vulnerability using a predator scent model of PTSD

**DOI:** 10.1101/438952

**Authors:** David Dopfel, Pablo D. Perez, Alexander Verbitsky, Hector Bravo-Rivera, Yuncong Ma, Gregory J. Quirk, Nanyin Zhang

## Abstract

Only a minority of individuals who experience traumatic event(s) subsequently develop post-traumatic stress disorder (PTSD). However, whether differences in vulnerability to PTSD result from predisposition or a consequence of trauma exposure remains unclear. A major challenge in differentiating these possibilities is that clinical studies focus on individuals already exposed to traumatic experiences, and do not take into account pre-trauma conditions. Here using the predator scent model of PTSD in rats and a longitudinal design, we measured pre-trauma brain-wide neural circuit functional connectivity (FC), behavioral and corticosterone responses to trauma exposure, and post-trauma anxiety. Individual differences in freezing responses to predator scent exposure correlated with differences in pre-trauma FC in a set of neural circuits, especially in olfactory and stress-related systems, indicating that pre-existing function in these circuits could predispose animals to differential fearful responses to threats. Counterintuitively, rats with the lowest freezing showed more avoidance of the predator scent, a prolonged corticosterone response, and higher anxiety long after exposure. This study provides a comprehensive framework of pre-existing circuit function that determines threat response strategy, which might be directly related to the development of PTSD-like behaviors.

## Introduction

Maladaptive response to trauma may lead to chronic stress disorders like posttraumatic stress disorder (PTSD). While trauma exposure is a feature of PTSD, it is not an unequivocal predictor of the disorder. In fact, only a minority of individuals who experience traumatic event(s) subsequently develop PTSD. However, mechanisms underlying this individual difference in vulnerability remain unclear.

It has been suggested that vulnerability to PTSD may result from individual differences in early life events, preexisting conditions, or responses to trauma exposure ^1-3^. However, mixed results have been reported. For instance, reduced hippocampal volume has long been considered a marker of PTSD. Nevertheless, some studies suggest that PTSD-related decreases in hippocampal volume might be a consequence of traumatic events and PTSD development, whereas other studies showed that PTSD patients’ twins who did not have a trauma experience also had smaller hippocampal volumes, suggesting that reduced hippocampal volume may pre-dispose individuals to developing PTSD ^4-7^. These conflicting results are also reported in studies of cortisol levels and brain activity in relation to PTSD ^8-10^, which together contribute to our elusive understanding of individual vulnerability to PTSD.

A major obstacle of investigating risk factors of PTSD is rooted in the difficulty of longitudinally monitoring of PTSD development pre-through post-trauma in humans with well-controlled traumatic events. Studies in humans focus on populations already exposed to a variety of uncontrolled traumatic events ^11^. Such barriers can be overcome in experimental animals, for which a longitudinal design with controlled traumatic stressors can be conveniently applied. Among many stress paradigms that have been used to produce rat models of PTSD, predator scent exposure in an inescapable environment is a commonly used psychological stressor ^12^. This method mimics a life-threatening situation that is ecologically relevant and physically innocuous, preventing confounding effects of inflammation or pain due to injury. Recent work with this model has found that PTSD-like behaviors are exhibited following exposure to predator scent ^13^. Importantly, Cohen et al. distinguished the responses of mal-vs. well-adapted rats, showing a 25-30% prevalence of PTSD-like behavior in rats exposed to trauma ^14^. This work highlights the potential for this animal model to investigate relevant individual variability in PTSD-like responses to trauma. Missing from previous studies, however, is information about preexisting conditions of animals that might predict their immediate and long-term responses to trauma.

Additionally, little is known about whether and how specific defense responses to a threat correlate with susceptibility to maladaptation. Exposure to a stressor elicits different fear responses depending on the perceived proximity and intensity of the threat. Moderate stressors produce passive freezing, while imminent predation threats elicit a fight or flight response ^15,16^. However, whether behavioral responses to predator scent is a useful indicator of subsequent development of PTSD is unknown.

Here we employed a longitudinal approach to studying individual variability of behavioral and functional outcomes of the predator scent animal model of PTSD. Resting-state functional magnetic resonance imaging (rsfMRI) data were collected from awake rodents before trauma exposure to record pre-existing neural circuit function using the awake rat rsfMRI approach established in our lab ^17-20^. Awake rat rsfMRI avoids confounding factors from anesthetics and permits correlation with behavior ^21-23^. We found that individual differences in behavior and neural circuit function prior to trauma exposure predicted animals’ susceptibility to developing PTSD-like behaviors. This suggests that pre-existing traits such as anterior olfactory nucleus (AON)-amygdala connectivity may be critical for determining individual resilience/susceptibility to developing PTSD.

## Results

The experimental schedule was summarized in Fig. 1a. Rats were first scanned using rsfMRI, then exposed to fox urine, and then given an anxiety test approximately one week after exposure.

**Figure 1.**
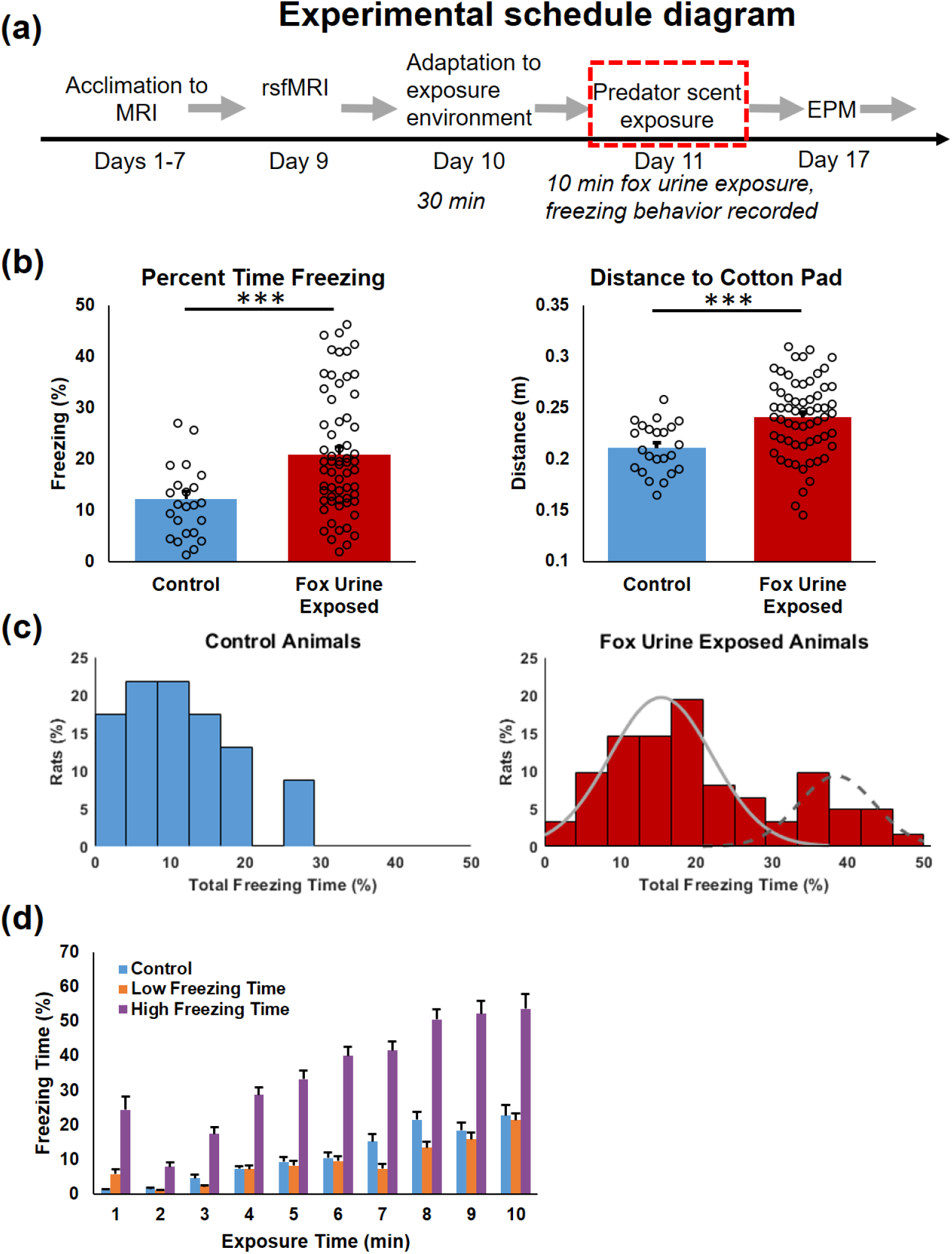
Freezing time during the 10-min exposure session in control (n = 23) and fox urine exposed (n = 62) groups. **a**) Diagram of experimental schedule. **b**) Fox urine exposed animals showed a significantly longer freezing time (left, p < 0.0005, in percentage) and greater mean distance (right, p < 0.0007, in meters) from the cotton pad than controls. **c**) The distributions of individual animals’ freezing time in the control group (left) and the exposed group (right) displayed as probability histograms. The freezing time distribution in exposed animals was fit to a Gaussian mixture model which demonstrated a bimodal distribution. **d**) Freezing time each min plotted as a function of time in low-freezing (n=21), high-freezing (n=21) and control animals during fox urine exposure. The high freezing animals displayed higher freezing phenotype during the first minute of exposure, with a significantly longer freezing time than both low freezing and control animals from the 1st minute through the 10th minute (p < 0.005 for each minute of exposure).

### Freezing responses to fox urine were highly variable across animals

A total of 87 Long-Evans (LE) rats were exposed to a predator scent stressor in the form of a compressed cotton pad sprayed with fox urine in an inescapable cage for 10 min. Freezing and location within the cage were quantified with behavioral tracking software (ANY-maze, Stoelting Co., Reston, VA). Compared to control animals that were exposed to a clean cotton pad, the group exposed to predator scent showed increased freezing (t-test, t_87_ = 3.660, p = 0.00044, Fig. 1b, left), and maintained a significantly greater distance from the cotton pad (t-test, t_87_ = 3.568, p = 0.00060, Fig. 1b, right). These data confirm that predator scent exposed animals displayed a significantly greater stress response compared to controls.

Closer examination of animals’ freezing time revealed distinct distributions between groups. Fig. 1c showed the histograms of freezing time in both groups. Control animals exhibited a unimodal distribution of freezing, whereas predator scent exposed animals displayed large inter-subject variability with a bimodal distribution (Fig. 1c, right panel) fitted with a mixed Gaussian distribution (bimodal Gaussian function provided the best fit relative to uni-modal and tri-modal Gaussian functions, model fit quality (the model with the lowest AIC is the best model to fit the data): Δ_1_ (AIC) = 11, Δ_2_ (AIC) = 0, Δ_3_ (AIC) = 6; Akaike weight for each model: *w*_1_ = 0.0039, *w*_2_ = 0.947, *w*_3_ = 0.0488). To further investigate this variation in freezing behavior, predator scent exposed animals in the top and bottom tertiles of freezing time were separated into high-freezing and low-freezing subgroups, respectively (n = 21 for each subgroup). Fig. 1d plotted freezing time as a function of exposure time in high-freezing, low-freezing and control groups. Interestingly, it was found that freezing differed between high-and low-freezing animals even during the first minute of predator scent exposure (two sample t-test, p< 0.009) and this difference became larger over the time course of exposure. There was no difference between low-freezing rats and controls. There was also no significant difference in mobility among all three groups, as evidenced by the maximum speed during exposure (one-way ANOVA, F_(2, 62)_ = 0.86, p = 0.43, Supplement Fig. 1).

### Low-, but not high-freezing animals were susceptible to developing PTSD-like behaviors

The fact that freezing time was almost identical between low-freezing and control animals might suggest that low-freezing animals had a similar stress response to controls. However, the spatial distribution of time spent in the cage during exposure suggested a different interpretation (Fig. 2). Control animals did not show a side preference with a near normal distribution of the time spent close to the center of the cage (Fig. 2a). In contrast, both high and low freezing animals exhibited a leftward skewed distribution of time spent, farther away from the fox urine. The average distance from the cotton pad was significantly greater in both low-and high-freezing animals compared to controls (One-way ANOVA, F_(2, 62)_ = 10.97, p = 0.000050 and p = 0.031, respectively), indicating avoidance behavior in exposed animals. This difference was also supported by the normalized freezing time at different distances from the cotton pad, as shown in Fig. 2b. Similar to the distribution of the total time spent in the cage, control animals did not seem to have any preference for the side of the cage in which they froze, whereas both high-and low-freezing animals tended to freeze farther away from the fox urine sample. Relative to high-freezing animals, low-freezing animals showed even more leftward bias. The normalized freezing time was significantly less in the zone 24 cm away from the pad (Two-way Repeated ANOVA, F_Group × Zone_ (22, 682) = 3.00, p = 0.0000059; Tukey-Kramer post-hoc test, low-vs high-freezing at 24 cm: p = 0.04), indicating that low-freezing animals were somewhat more avoidant than high-freezing animals. This can also be seen in the heat plots of Fig. 2a. Prior to predator scent exposure during the habituation period, the three groups showed no significant differences in freezing time (One-way ANOVA, F_Group × Zone_ (2, 62) = 0.24, p = 0.78, Supplement Fig. 2) or location distribution (the cage was divided into three even-size zones, Two-way ANOVA, F_Group × Zone_ (4, 186) = 1.28, p = 0.28).

**Figure 2.**
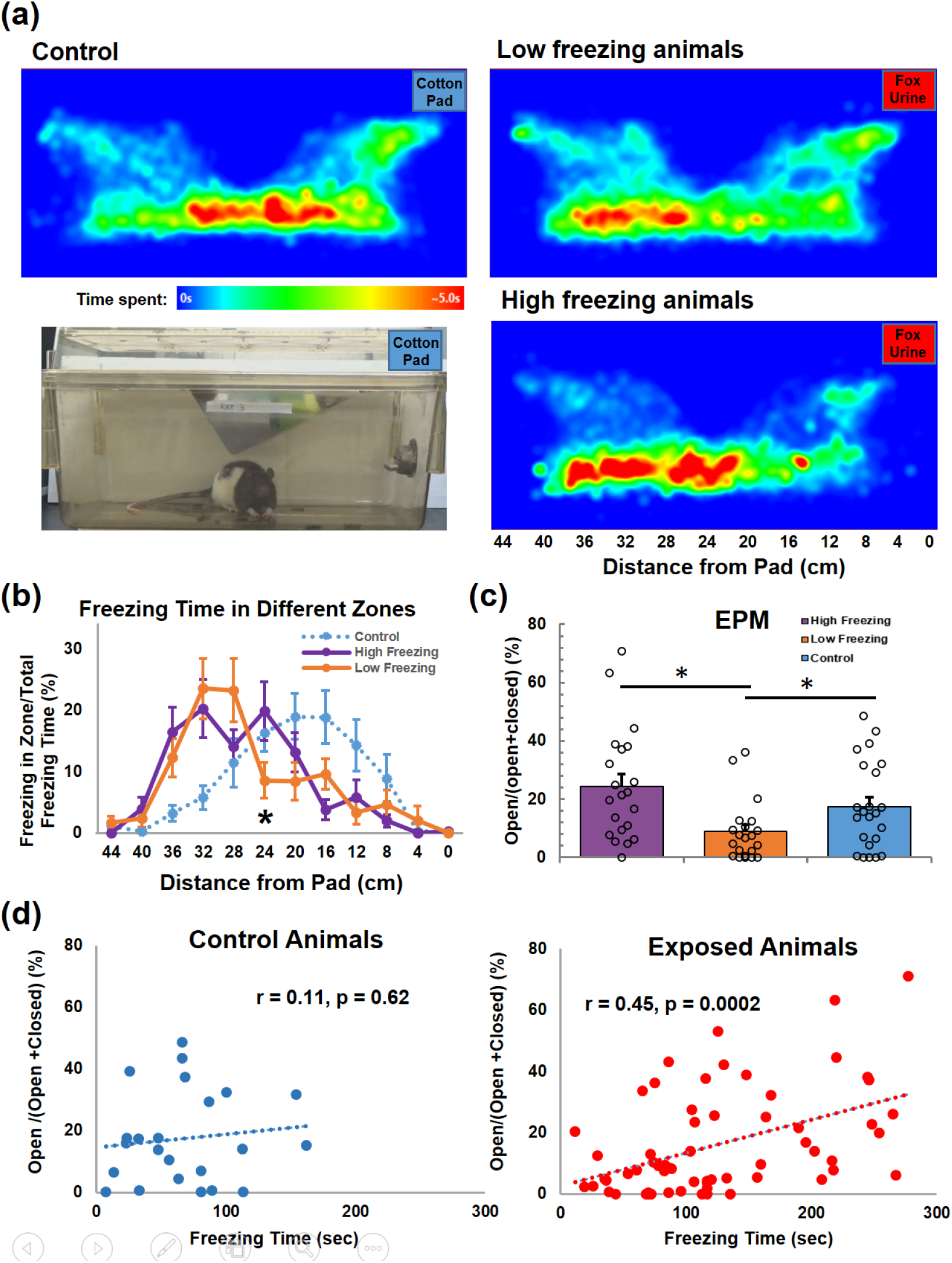
Low-freezing animals exhibited stronger stress response during fox urine exposure and higher anxiety 6 days after exposure. **a**) Heat maps of the total time at each location for the control, low freezing, and high freezing groups. The control group showed no evident bias for either side of the cage. The low freezing group showed a strong bias toward the far end (from the pad) of the cage, while the high freezing rats also showed a bias toward the far end of the cage, but to a lesser extent. **b**) The distribution of the percent freezing time for each group across the cage, divided into 12 equivalent-sized zones. The x-axis indicates the distance to the pad from the center of each zone. Similar to the distributions of total time spent, the percent freezing time in control rats displayed little bias for either side of the cage. High and low freezing animals showed a bias of percent freezing time toward the far end of the cage, with the low-freezing rats showing even stronger bias than high-freezing animals, evidenced by significantly lower percent freezing time of the low-freezing group in the zone 24 cm away from the pad compared to the high-freezing group (p =0.026). **c**) Elevated plus maze performed 6 days post exposure in all three groups of animals. Low-freezing animals showed a significantly lower EPM score (open/open+closed arm time) compared to both control (p = 0.030) and high-freezing animals (p = 0.0017), while no significant difference was seen between control and high freezing groups. **d**) The freezing time was positively correlated to the EPM score across all exposed animals (r = 0.45, p = 0.0002), but not across control animals (r = 0.11, p = 0.62).

To assess the long-lasting PTSD-like behaviors in exposed animals, we measured anxiety levels using an elevated-plus maze (EPM) 6 days after predator scent exposure. Resembling the observation of increased avoidance, low-freezing animals exhibited heightened long-lasting anxiety compared to both control (t-test, t = 2.24, p = 0.030) and high-freezing animals (t-test, t = 3.37, p = 0.0017), as indicated by the ratio of open/open+close arm time in EPM tests (Fig. 2c). There was no statistical difference between high-freezing and control animals, indicating that low-, but not high-freezing animals were susceptible to developing PTSD-like behaviors following exposure to predator scent. In addition, freezing was positively correlated with the EPM score in exposed animals (R^2^ = 0.21, p = 0.0002), but not controls (R^2^ = 0.01, p = 0.62, Fig. 2d). These results demonstrate that behavioral responses during trauma exposure can predict long-term maladaptation. Furthermore, in a separate experiment (Supplement Fig. 3) low-freezing animals were also associated with other types of PTSD-like behaviors including more marble burying behavior (n = 5 each group, two sample t-test, t = 5.47, p = 0.048), less habituation to acoustic startle response (n = 9 each group, two-sample t-test, t = 5.3, p = 0.050) and less pre-pulse inhibition (n = 9 each group, two-sample t-test, t = 5.16, p = 0.037) than high-freezing animals 6 days post predator scent exposure. This provides additional evidence that low-freezers were vulnerable to developing PTSD-like behaviors.

### Low-freezing animals showed prolonged corticosterone response to predator scent

We also measured the corticosterone (CORT) level immediately before predator scent exposure (0 min, baseline) as well as 30 min, 60 min and 120 min after exposure (Fig. 3, 15 controls including 9 exposed to air and 6 exposed to lemon scent, as well as 24 fox urine exposed animals). Again, we grouped rats according to their freezing time during predator scent exposure with 12 low-freezing and 12 high-freezing animals (we did not use tertiles given a relatively smaller sample size in this experiment).

**Figure 3.**
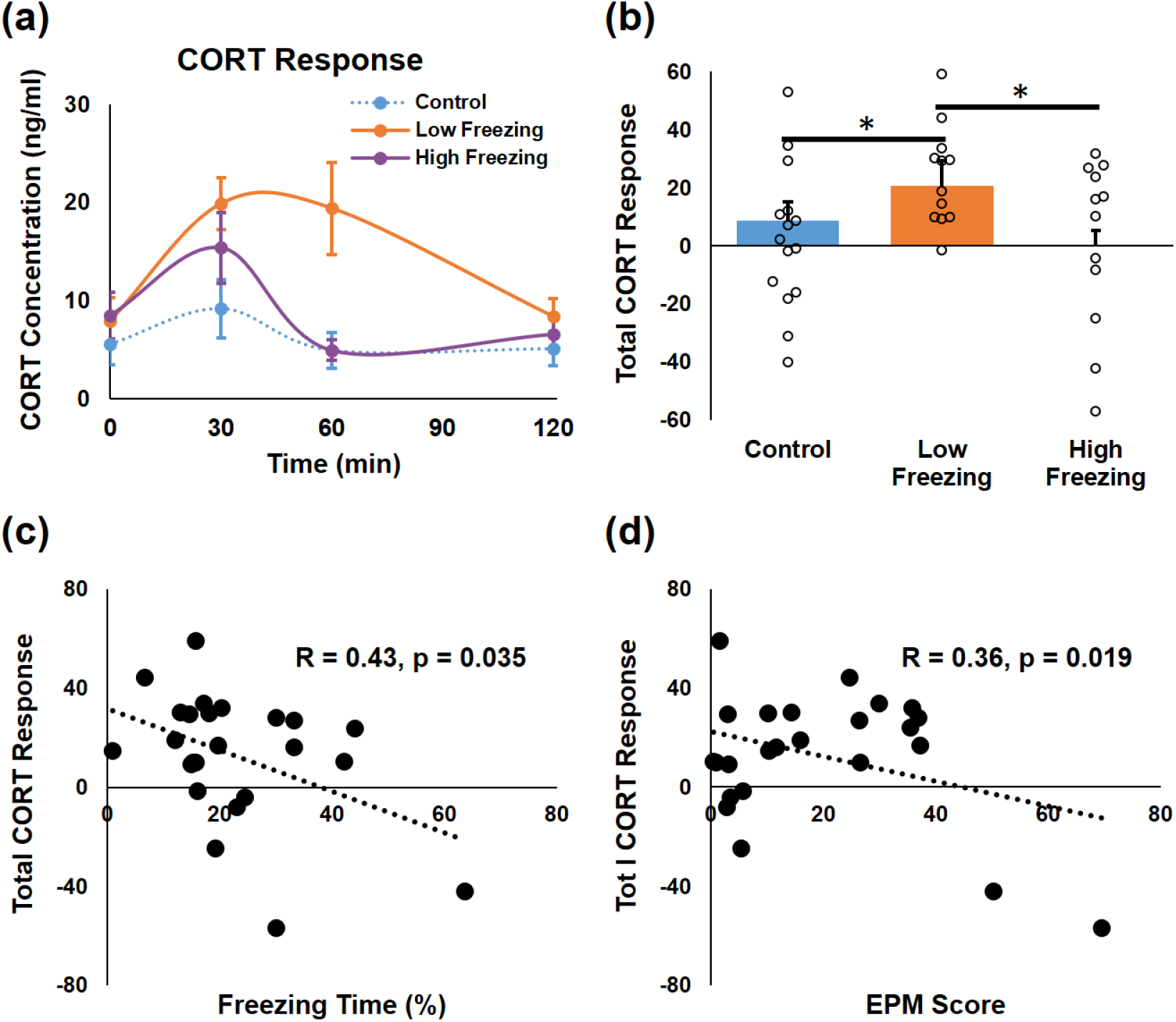
Corticosterone (CORT) response to fox urine exposure. **a**) The time course of CORT (in ng/ml) for high-, low-freezing and control animals. There was a significant difference in CORT level changes over time among three groups (Two-way repeated ANOVA, F_group × time_ (6,108) = 2.84, p = 0.023). Control animals showed no appreciable CORT change across time (One-way repeated ANOVA, F_time_ (3,56) = 0.8, p = 0.50). High-freezing animals exhibited a relatively short CORT response, peaking at 30 min and returning to baseline at 60 min (Tukey-Kramer test, 30 min vs 60 min: p = 0.049; 0 min vs 60 min: p = 0.54; 60 min vs 120 min: p = 0.93). The peak amplitude of CORT response at 30 min was comparable between high-and low-freezing animals (Tukey-Kramer test, p = 0.59). However, relative to high-freezing rats, the CORT response in low-freezing animals was prolonged, maintained at a high level at 60 min (Tukey-Kramer test, 30 min vs 60 min: p = 1; 0 min vs 60 min: p = 0.0005) and had a delayed return to baseline at 120 min (Tukey-Kramer test, 60 min vs 120 min: p = 0.0098; 0 min vs 120 min: p = 1). In addition, low freezers’ CORT level was significantly higher at 60 min than high freezers (Tukey-Kramer test, p = 0.0039). **b**) Total CORT response was quantified by the area-under-the-curve (AUC) of the CORT response curve, with the baseline CORT level (i.e. 0 min) being subtracted from each time point. This quantification method takes into account both the amplitude and length of the hormone response and represents the total output of the hypothalamus-pituitary-adrenal axis during exposure. Low-freezing animals showed a significant higher total CORT response than high-freezing (p = 0.031) and control (p = 0.017) animals. **c**) Significant correlation between the total CORT responsive (y-axis) and freezing time during exposure across all fox urine exposed animals (r = 0.43, p = 0.035). **d**) Significant correlation (r = 0.36, p = 0.019) between the total CORT response (y-axis) and EPM score (open/open+closed arm time, x-axis) measured 6 days post exposure across all animals tested.

There was a significant difference in CORT level changes over time among groups (Two-way repeated ANOVA, F_group × time_ (6,108) = 2.84, p = 0.023, Fig. 3a). In particular, control animals showed no significant CORT change across time (One-way repeated ANOVA, F_time_ (3,56) = 0.8, p = 0.50). High-freezing animals exhibited a relatively short CORT response, peaking at 30 min and returning to baseline at 60 min (Tukey-Kramer test, 30 min vs 60 min: p = 0.049; 0 min vs 60 min: p = 0.54; 60 min vs 120 min: p = 0.93). The peak amplitude of CORT response at 30 min was comparable between high-and low-freezing animals (Tukey-Kramer test, p = 0.59). However, relative to high-freezing rats, the CORT response in low-freezing animals was prolonged, maintained at a high level at 60 min (Tukey-Kramer test, 30 min vs 60 min: p = 1; 0 min vs 60 min: p = 0.0005) and had a delayed return to baseline at 120 min (Tukey-Kramer test, 60 min vs 120 min: p = 0.0098; 0 min vs 120 min: p = 1). In addition, low freezers’ CORT level was significantly higher at 60 min than high freezers (Tukey-Kramer test, p = 0.0039).

The total CORT response was quantified by the area-under-the-curve (AUC) of the CORT response curve with the baseline CORT level (i.e. 0 min) being subtracted from each time point. This quantification method took into account both the amplitude and duration of the hormone response. Low freezers showed significantly higher total CORT response than high freezers (t = 5.29, p = 0.031, Fig. 3b), as well as controls (t = 6.58, p = 0.017, Fig. 3b). Furthermore, the total CORT response was significantly correlated with freezing time across all fox urine exposed animals (R = 0.433, p = 0.035, n = 24, Fig. 3c), and was also linked to long-lasting anxiety, evidenced by a significant correlation with the EPM score measured 6 days after exposure across all animals tested (r = 0.36, p = 0.019, n= 39, Fig. 3d). Taken together, these data demonstrate that low-freezing animals displayed prolonged CORT response to predator scent stress, which was related to long-term anxiety.

### Pre-trauma neural circuit functional connectivity predicted fear responses

In addition to behavioral tests, rsfMRI was performed before trauma exposure to measure preexisting resting-state functional connectivity (RSFC) in neural circuits across the whole brain. Echoing our behavioral data showing difference in freezing during the very beginning of predator scent exposure, RSFC of multiple widespread neural circuits significantly differed between high-and low-freezing animals (false discovery rate (FDR) = 0.01, Supplement Fig. 4a). These regions/circuits have been implicated in stress responses, processing of emotionally valent stimuli, olfactory function and freezing behavior.

To identify the specific neural circuits correlated with the freezing phenotype, we conducted a data-driven correlational analysis for all neural connections (67 regions covering the whole brain, 2211 connections in total) between their RSFC and the cumulative freezing time across all exposed animals (Fig. 4a, upper triangle). 15 neural circuits exhibited a significant correlation between pre-exposure RSFC and freezing time (p < 0.05, FDR corrected, Fig. 4a, lower triangle). Relative to high-freezing animals, six of these connections showed stronger connectivity, whereas 9 showed lower connectivity in low-freezing animals, suggesting that the pre-trauma connectivity within these circuits might have predictive value for the freezing behavior during exposure. Fig. 4b showed three representative RSFC maps, respectively generated with one region in these 15 connections as the seed (i.e. anterior cingulate cortex, AON and parabrachial nucleus). Fig. 5 showed the correlational relationship between pre-trauma RSFC and freezing time for all 15 circuits. The diagram of this freezing-associated network was summarized in Fig. 6.

**Figure 4.**
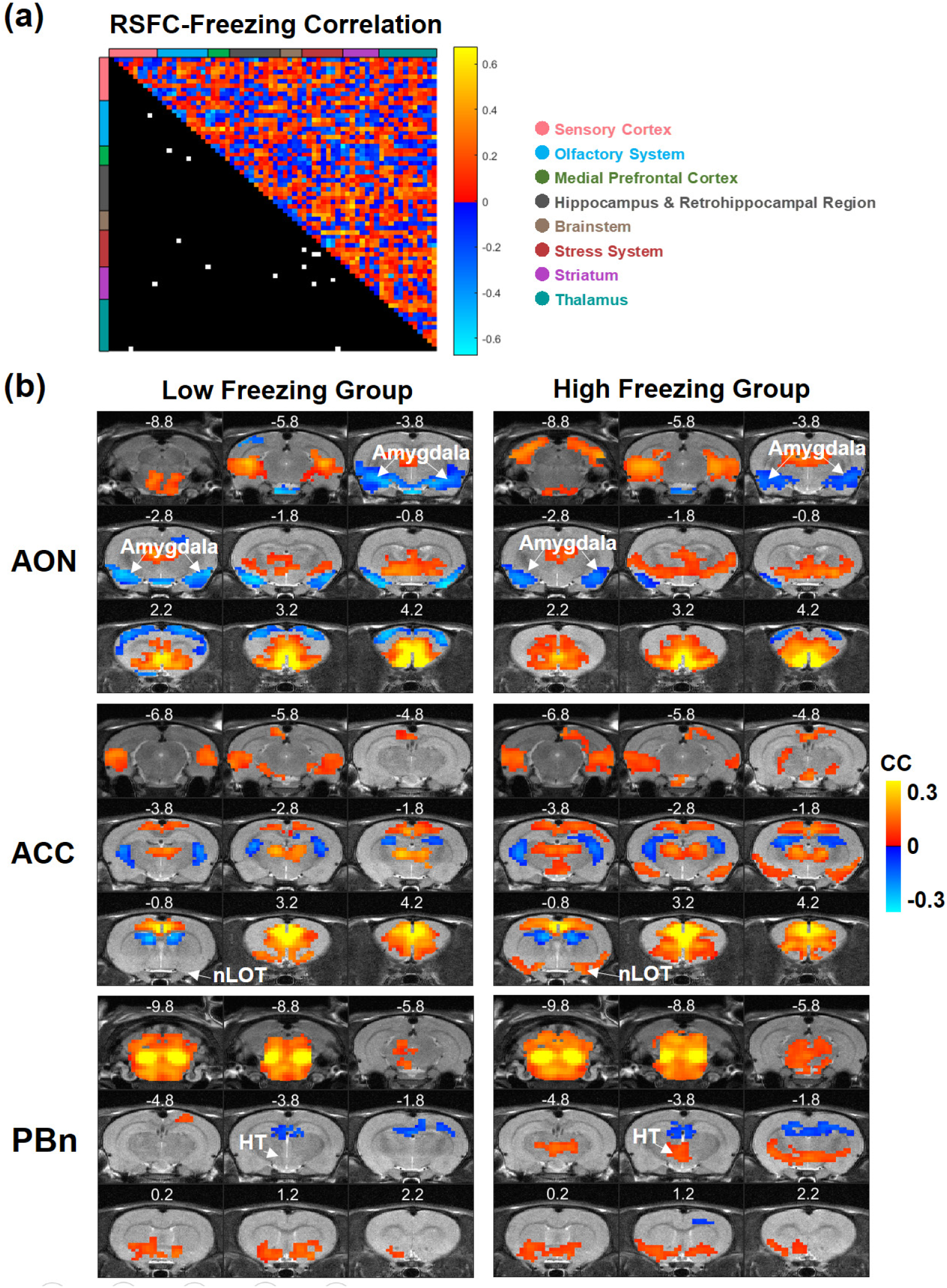
**a**) Correlations between pre-trauma RSFC and freezing time for all connections (upper triangle) and super-threshold connections (lower triangle, each white element represents a super-threshold connection). Brain regions are arranged based on the brain system (color coded) they belong to. **b**) Three representative RSFC maps, generated with ACC, AON and PBn as the seed, respectively (colored voxels, p < 0.05, FDR corrected). Distance to Bregma was labeled at the top of each slice. ACC, anterior cingulate cortex; AON, anterior olfactory nucleus; HT, hypothalamus; nLOT, lateral olfactory tract nucleus; PBn, parabrachial nucleus.

**Figure 5.**
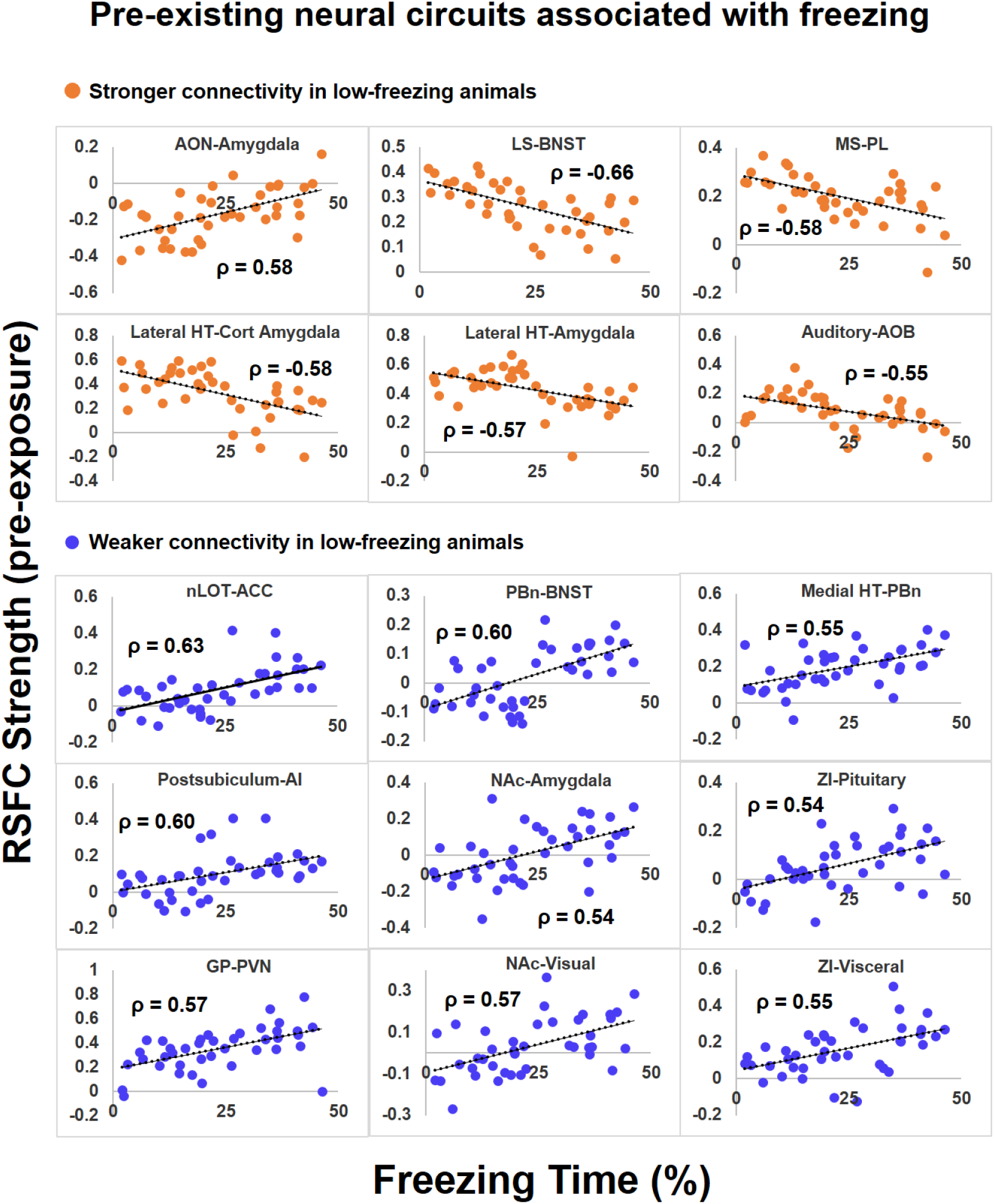
Neural circuits with pre-trauma resting-state functional connectivity (y-axis) correlated (p<0.05, FDR corrected) with percentage freezing time (x-axis) in all fox urine exposed animals (n=62). AON, anterior olfactory nucleus; LS, lateral septum; BNST, bed nucleus of the stria terminalis; MS, medial septum; PL, prelimbic cortex; HT, hypothalamus; Cort Amygdala, cortical amygdala; AOB, accessory olfactory Bulb; nLOT, lateral olfactory tract nucleus; ACC, anterior cingulate cortex; PBn, parabrachial nucleus; AI, postsubiculum-agranular insula; NAc, nucleus accumbens; ZI, zona inserta; GP, globus pallidus; PVN, paraventricular nucleus.

**Figure 6.**
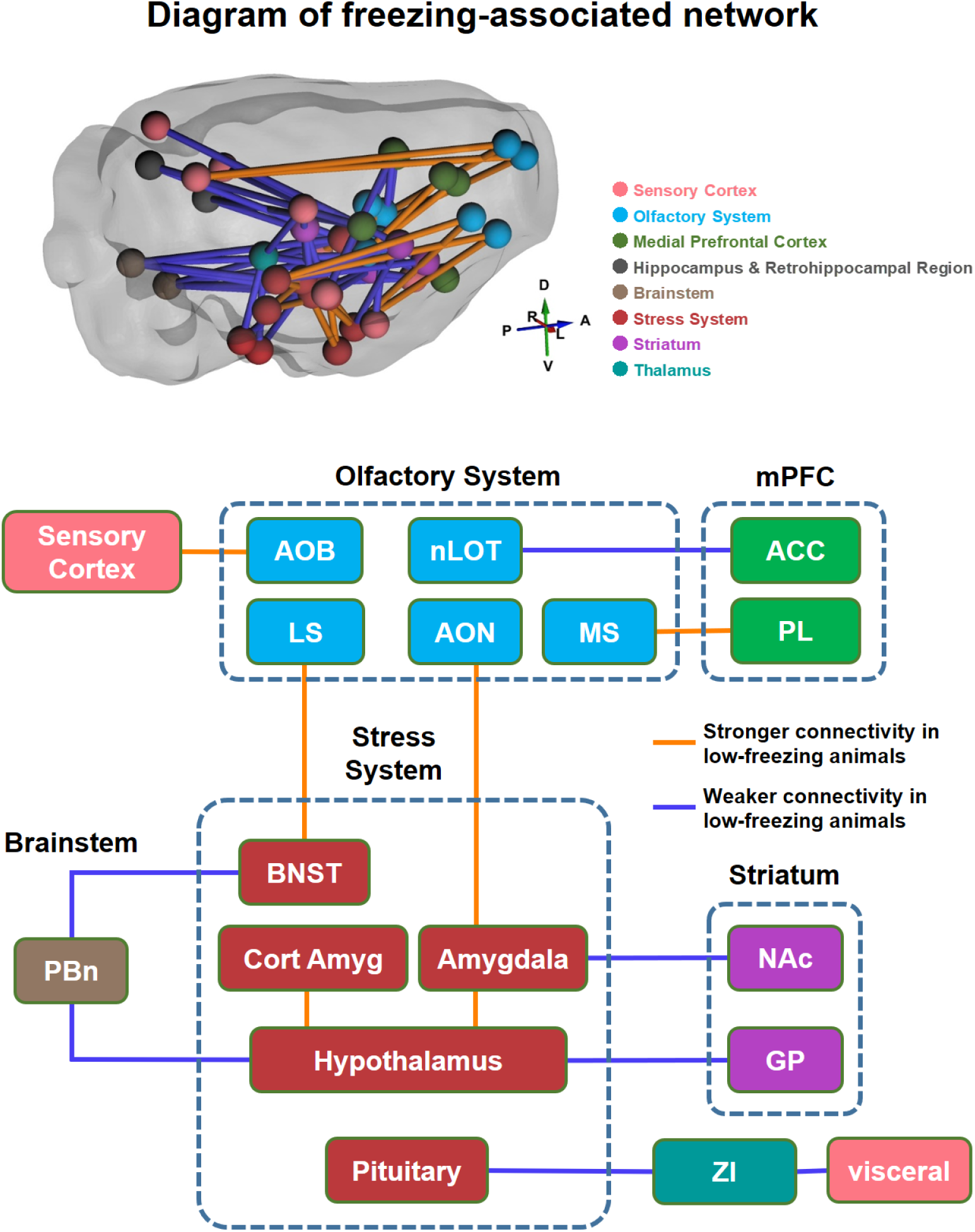
Diagram of freezing-associated neural network, including neural circuits showing significant correlation between pre-trauma RSFC and total freezing time shown in Fig. 5. Brain regions are grouped and color coded based on the brain systems they belong to. Orange lines indicate stronger connectivity in low-freezing animals. Blue lines indicate weaker connectivity in low-freezing animals. See the caption of Fig. 5 for the abbreviation.

A closer examination of these 15 circuits revealed a comprehensive network particularly involving the olfactory and stress-related brain areas. Specifically, low-freezing animals showed stronger pre-trauma connectivity in two olfactory circuits (i.e. AON-amygdala, and olfactory bulb-auditory cortex), two septal circuits (i.e. lateral septum (LS)-bed nucleus of the stria terminalis (BNST), medial septum (MS)-prelimbic cortex (PL)), and two circuits within the stress system (lateral hypothalamus-cortical amygdala and lateral hypothalamus-amygdala). Stronger connectivity in the two olfactory circuits suggest that odor may exert more influence on the amygdala and cortex in low-freezing animals. Additionally, the LS-BNST projection might underlie stronger predator scent-induced stress response in low-freezing animals. In support of this, both the LS and BNST are activated by predator odor ^24,25^, and the BNST is actively involved in sustained fear response to diffuse threat ^26^. Stronger functional connectivity within the stress system can also contribute to higher stress response and anxiety in low freezing animals.

Conversely, nine neural circuits showed weaker pre-trauma functional connectivity in low-freezing animals, including two parabrachial nucleus (PBn) circuits (PBn-BNST and PBn-hypothalamus), two nucleus accumbens (NAc) circuits (NAc-amygdala and NAc-visual cortex), two zona inserta (ZI) circuits (ZI-pituitary gland and ZI-visceral cortex), as well as the circuits of lateral olfactory tract nucleus (nLOT)-anterior cingulate cortex (ACC), postsubiculum-agranular insula (AI), and globus pallidus (GP)-paraventricular (PVN) hypothalamus. PBn is tightly related to olfactory function and electrophysiological data show that calcitonin gene–related peptide (CGRP) in PBn has inhibitory effects in the BNST ^27^. Therefore, weaker connectivity in the PBn-BNST circuit suggests less inhibition in BNST and could contribute to heightened anxiety in low freezing animals. Weaker RSFC between the nLOT and ACC can reduce ACC’s control over predator scent-triggered defense reaction in low-freezing animals. In addition, the NAc-amygdala circuit mediates avoidance behaviors ^28^, and decreased NAc connectivity was consistent with report of the inverse association between NAc activity and the intensity of emotional stress response ^24^, which might also be related to goal-directed motor behavior ^29^. Furthermore, lower connectivity in ZI circuits (i.e. ZI-pituitary gland and ZI-visceral cortex) may be involved in control of sympathetic visceral reactions and the hypothalamus–pituitary–adrenal (HPA) axis activity ^30^. Lastly, the GP-PVN hypothalamus projection, known to involve corticotropin-releasing factor (CRF) neurons, were reported to link stress response to motor output ^31^.

Taken together, these data indicate that differences in functional connectivity in neural circuits particularly involving the olfactory and stress-related systems might predispose animals to differential stress responses during predator scent exposure, which can be linked to individual difference in subsequent development of PTSD-like behaviors long after trauma exposure.

### Fear responses and long-term anxiety in fox-urine exposed animals were not due to the novelty of the fox urine scent

To rule out the possibility that differential responses in fox-urine exposed animals resulted from the novelty component of the scent, we repeated the experiment with a lemon scent being used as the control scent in a separate cohort of animals (n=24). Lemon exposed animals showed virtually identical freezing (Supplement Fig. 5a, left, n = 6, two-sample t-test, t = 2.0, p = 0.16) and avoidance (distance from the pad, Supplement Fig. 5a, right, two-sample t-test, t = 1.14, p = 0.32) compared to animals exposed to no scent (i.e. air, n = 6), which ruled out the possibility that the novelty nature in the scent affected fear responses during exposure. In contrast, the fox urine exposed group (n = 12) maintained a greater mean distance from the cotton pad (one way ANOVA, F_(2,23)_ = 11.61, p = 0.0004) than both air (p = 0.012) and lemon (p = 0.0006) exposed animals. In addition, fox urine exposed animals showed a trend of more freezing than non-traumatized animals (left, p = 0.056, lemon and air groups combined). Furthermore, consistent with the findings in the main cohort of animals, low-freezing animals exhibited stronger stress response during fox urine exposure and higher anxiety 6 days after exposure. Supplement Fig. 5b showed heat maps of the spatial distribution of time spent in the cage for the air-exposed, lemon-exposed, fox urine-exposed low freezing, and fox urine-exposed high freezing groups, respectively. The air and lemon groups showed no evident bias for either side of the cage, whereas the low-freezing group showed a strong bias toward the far end (from the pad) of the cage. High freezing rats also showed a bias toward the far end of the cage, but to a lesser extent. This result was also confirmed in the spatial distribution of the normalized freezing time for each group (supplement Fig. 5c), which displayed virtually no spatial bias in air and lemon groups, but leftward skewed distributions in high and low freezing rats, with an even stronger bias in low-freezing rats. Supplement Fig. 5d showed the long-term anxiety measured using EPM performed 6 days post exposure in all four groups of animals. Low freezing animals showed a significantly lower EPM score (one way ANOVA, F_(2,17)_ = 4.2, p = 0.033) compared to high freezing animals (p = 0.026). Also consistent with our data in the main cohort of animals, freezing time was positively correlated to the EPM score across all fox-urine exposed animals (r = 0.783, p = 0.0026, Supplement Fig. 5e), but not in non-traumatized animals (r = 0.213, p = 0.506, Supplement Fig. 5e). Taken together, these results ruled out the possibility that differential short-term and long-term behavioral responses in fox-urine exposed animals was due to the novel nature of the fox urine scent.

### Differential fear behavior during exposure and long-term anxiety between low-and high-freezing animals were not caused by pre-exposure imaging

To rule out the possibility that differential fear response during predator scent exposure and long-term anxiety between low-versus high-freezing animals indeed resulted from predator scent exposure as oppose to other factors such as the acclimation and/or imaging procedures, we also divided control animals into tertiles based on their cumulative freezing time. No differences in avoidance behaviors including distance to the cotton pad (two-sample t-test, t = 1.22, p = 0.25), the distribution of time spent in the cage (two-way ANOVA, F_2,28_ = 2.20, p = 0.13, also see Supplement Fig. 6a), the distribution of freezing time in the cage (repeated measures ANOVA, F_2,154_ = 0.34, p = 0.98), or long-term anxiety level (EPM score, two-sample t-test, t = 0.37, p = 0.72) were observed between low-and high-freezing control animals. In addition, unlike predator scent-exposed animals, no neural connections exhibited correlation between pre-exposure RSFC and freezing time, and no connection displayed significant RSFC difference between low-and high-freezing control animals (Supplement Fig. 4b), indicating that these animals showed virtually no difference in neural circuit function.

Furthermore, we repeated the fox urine exposure experiment in a separate cohort of animals that were not previously acclimated or imaged (control, n = 20, fox urine exposed, n = 35). Similar differential response during exposure was found in these animals, with lower freezers generally exhibiting more avoidance from the fox urine (Supplement Fig. 6b). These data suggest that the inherent difference between low and high freezers was not caused by potential differential responsiveness to acclimation. This result was supported by the data showing that all three groups of animals (i.e. low-freezing, high-freezing and control) in the main cohort were acclimated to a similar degree, reflected by their consistent motion levels during imaging (One-way ANOVA, F_(2,62)_ = 1.23, p = 0.3, Supplement Fig.4c). These findings are also consistent with our previous publications demonstrating that acclimation/imaging does not mask, nor does it interact with the effect of predator stressor ^21,23,32^. Taken together, these results confirmed that differential fear behaviors and anxiety levels between low-and high-freezing animals were not driven by the acclimation/imaging procedures as all control animals underwent the identical procedures including imaging, except for being exposed to predator scent.

## Discussion

Employing a longitudinal design, we investigated animals’ individual variability in stress response and subsequent development of PTSD-like behaviors following predator scent exposure. We found that animals displayed large differences in freezing behavior during fox urine exposure. Animals expressing less freezing exhibited increased avoidance and tended to develop long-lasting heightened anxiety. Importantly, this difference in behavioral responses was apparent at the beginning of predator scent exposure, suggesting pre-existing vulnerability. This notion was supported by the rsfMRI data showing significant pre-trauma difference in neural circuit connectivity between high-and low-freezing animals. In addition, low-freezing animals showed a prolonged CORT response to predator scent, which was correlated with freezing and long-lasting EPM anxiety measurement. Taken together, this study provides important evidence supporting the hypothesis that preexisting circuit function determines the defense response strategy during threat and that this behavioral response might be directly related to the development of PTSD. It also highlights the importance of studies that analyze factors both before and after trauma for the study of PTSD.

### Freezing time alone might not be a reliable measure of response to threat

Behavioral response to stress is commonly categorized into fight, flight or freezing. In rodent behavior testing, freezing is dominantly used as the measure of threat reaction. Although freezing is a universal fear response, it manifests at intermediate levels of predator threat ^16^. Meanwhile, other types of threat-induced responses including fight-or-flight, which is elicited during imminent threat ^15^, and avoidance can counteract the measure of freezing. For this reason, the use of freezing as the only measure for stress response of an individual animal can be problematic if alternative coping strategies are not considered ^33^. Indeed, there is growing evidence suggesting the ambiguity of freezing time as it relates to stress ^15,34^.

In line with this concept, our data show that animals with the lowest freezing time are in fact the most stressed subgroup. Their highly biased distribution in the total time and freezing time at different distance to the cotton pad indicate that low-freezing animals might have perceived a higher threat level and used strategies other than freezing (e.g. avoidance), particularly at closer proximity to the pad. This result was corroborated by significantly higher total CORT response during exposure in low-freezing animals, relative to both high-freezing and control animals. This higher perceived threat level is likely a crucial factor in maladaptation to stress response (e.g. higher long-lasting anxiety level), which is in line with the literature investigating the role of threat perception in human PTSD diagnoses ^35^. Taken together, these results demonstrate that freezing time alone might not be a reliable measure of reaction to threat. These data also highlight the importance of separating subpopulations, in particular in stress studies, as key information can be lost through the averaging of subpopulations that display bidirectional responses.

It has to be noted that lower freezing behavior in low freezers did not reflect a reduced ability to respond to the predator scent. Low-freezing animals exhibited more locomotor activity, measured by distance travelled during exposure (Supplemental Fig. 1, One-way ANOVA, F_(2,62)_ = 8.43, p = 0.001, high freezers vs low-freezers, p = 0.001), and more active avoidance from the fox urine. Furthermore, during exposure there was no difference in mobility among low-, high-freezing and control groups, reflected by the maximal speed (Supplement Fig. 1), also suggesting that low freezers were not suppressed in their response to the fox scent. Different responses during exposure were also not originated from any systematic bias in these measures among animals, because during the habituation period prior to exposure, the three groups showed no difference in freezing time (Supplement Fig. 2) or location distribution. Thus, the difference in freezing to the predator scent likely reflects a difference in the defensive strategy selected.

### Behavioral and neuroendocrine reactions to a threat predict susceptibility to long-term maladaptation

Whether defensive reactions predict long-term emotion regulation and/or symptom development has been an unresolved issue, largely due to lack of longitudinal studies that follow up the relationship between behavioral reactions during trauma exposure and the development of psychopathology ^15^. Our longitudinal study demonstrates that freezing during trauma exposure predicts long-term maladaptation in animals, supporting that specific behavioral responses to a threat might predict susceptibility to long-term PTSD-like behaviors.

The relationship between the behavioral reaction during stress and vulnerability to stress maladaptation is corroborated by neuroendocrine measures during and following predator scent stress. We observed that in low-freezing (i.e. vulnerable) animals, the CORT response was prolonged and had a delayed return back to baseline after stress, and this pattern of CORT response has been suggested as a marker of maladaptation to stress in PTSD ^36,37^. Correspondingly, in our study, both freezing time and the total CORT response were correlated with the long-term anxiety measure. These data are also consistent with the theory that stress hormone production is an important and protective part of acute stress response triggering negative feedback on CORT production ^38,39,41,42^. In fact, it has been shown that supplementing CORT at the time of trauma ^44^, or immediately after trauma ^45^ has protective effects against PTSD symptom development ^38^. Taken together, our data indicate that individual variation in behavioral and hormonal responses to stress can be key predictors of the development of emotion regulation and chronic pathological states.

### Pre-trauma neural circuit function predisposes animals to differential stress response

#### i) Connections between the olfactory and stress systems

Our rsfMRI data suggest that sensory inputs play an important role in the stress response and subsequent development of PTSD-like behaviors. AON has been shown to be activated by variable stressors ^46^ including predator scent ^47^. The mitral/tufted (MT) cells in the olfactory bulb send their processes to AON ^48^, which projects to amygdala ^49^. This connectivity suggests that the flow of olfactory information through the AON to the amygdala could drive the stress response to predator scent. Stronger AON-amygdala connectivity in low-freezing animals suggest that the olfactory cue could have a stronger impact on amygdala activity, which would lead to stronger stress response. Conversely, weaker nLOT-ACC connectivity in low-freezing animals indicates less ACC control during predator scent exposure, which could affect the switching between freezing and active defense modes ^50^. These data also agree with the report that the olfactory cortex and its connections with the stress system are tightly linked to stress hormone responses to predator scent in rodents ^51^. Taken together, these results highlight the importance of sensory input in the stress response during predator scent exposure, which also suggest that weaker sensory drive in the stress system may help promote resilience to trauma.

In addition to stronger sensory-stress connectivity, stronger functional connectivity within the stress system was observed in low-freezing animals, consistent with their higher stress response and anxiety. In particular, it is known that threat-induced autonomic sympathetic responses are mediated by amygdala projections to the lateral hypothalamus ^15^. Stronger connectivity in this circuit could result in higher arousal, higher cardiovascular activity and increased muscle tone during predator odor exposure in low-freezing animals ^52^.

#### ii) BNST circuits

Our data also suggest that BNST may be involved in the neural network associated with differential stress response during predator scent exposure. BNST, as a part of the extended amygdala, is well known to be involved in sustained anxiety when a threat is diffuse and/or uncertain ^26,53^ such as predator odor ^25^. As a result, this region is particularly associated with anxiety ^54-56^. The anatomical connections between BNST and LS and PBn have been well documented ^27,57,58^, but the specific function of these projections is less well studied. It has been shown that CGRP in the PBn inhibits neurons of the BNST ^27^, and BNST projections to the PBn dampen anxiety response ^59^. These data support the idea that a weaker PBn-BNST connection in low-freezing (i.e. vulnerable) animals caused disinhibition of BNST activity, which led to a stronger stress response and heightened long-lasting anxiety. Conversely, LS mediated activity, possibly again driven by olfactory inputs ^24^, can potentiate BNST activity in these animals, and generate stronger stress responses and heightened anxiety.

#### iii) The hypothalamus-GP circuit

Another interesting circuit that is coupled to freezing behavior is the PVN-GP connection. CRF neurons in the PVN hypothalamus, amygdala and BNST send synapses to the external GP ^31^, which expresses high levels of the primary CRF receptor (i.e. CRFR1). These connections makes GP an entry point connecting the stress system with the basal ganglia, which links the stress-relevant information to the directed movement ^31^. Consistent with this notion, our data suggest that the pre-trauma connectivity in this circuit is directly associated with stress-related behaviors in animals.

#### iv) Possible contributions of other neural circuits to variable freezing behavior

Stronger connectivity between the MS and PL can be linked to stronger fear in low-freezing animals, given that PL promotes fear expression and avoidance ^60,61^. Also considering septal neurons are known to mediate the generation of theta oscillations ^62,63^, the MS-PL connection may contribute to fear-associated theta rhythm in prefronto-amygdalo-hippocampal pathways ^64^. In addition, the postsubiculum-insula circuit connectivity could be responsible for differential spatial movement in separate subgroups of animals.

Lastly, a number of additional connections showed significant difference in pre-trauma RSFC between low-freezing and high-freezing animals, even though their RSFC was not correlated with freezing time. For example, lower RSFC between CA3 and reticular nucleus (RT) was observed in low-freezing rats. RT is the hub connecting hippocampus with PFC, and reduced connectivity in vulnerable rats might be due to less prefrontal control of hippocampus ^65^. Although premature, these results suggest that preexisting neural circuit differences in vulnerable animals might reflect a sophisticated network, and PTSD should be viewed as a network phenomenon.

### Relevance to human PTSD

Our neural circuitry and neuroendocrine findings in animals echo PTSD research in humans. Altered sensory function has also been reported in PTSD patients ^66^. In addition, a significant number of brain regions identified here (e.g. BNST, PFC) are tightly related to anxiety disorders in humans ^67^, although the present study goes beyond static neuroanatomy of individual brain regions and focuses on pre-existing neural circuit function. Furthermore, human data also suggest that a dysregulated CORT response is a predictor of vulnerability to subsequent development of PTSD ^41,42,45,68^. Taken together, these results provide compelling evidence supporting the relevance of the present study to human PTSD.

In summary, our data demonstrate that differences in neural circuit function present before trauma exposure may predispose animals to specific stress responses and long-term PTSD-like behaviors. Future studies that focus on the causal relationship between the baseline neural circuit function and behavioral responses during trauma exposure and long-term outcome in PTSD-like behaviors will be needed to understand pathological susceptibility to PTSD. For instance, it will be of particular interest to test whether optogenetic manipulation of specific pathways based on the rsfMRI data during trauma exposure can alter stress response and development of long-term PTSD-like behaviors in animals.

## Methods

Adult male LE rats (250-350g) were obtained from Charles River Laboratories (Raleigh, North Carolina site). Rats were housed in pairs in Plexiglas cages with controlled ambient temperature (22-24 °C) on a 12/12 hour day-dark schedule. Food and water were provided ad libitum. A total of 87 rats were used in the present study, in which 23 rats were controls and 64 rats were exposed to predator scent using a single-episode predator scent exposure in an inescapable environment paradigm. Two rats were removed from the exposed group before analysis due to health concerns. All rats were used for behavior tests and rsfMRI experiments. All animal procedures were reviewed and approved by the Institutional Animal Care and Use Committee (IACUC) of the Pennsylvania State University. The experimental procedure was summarized in Fig. 1a.

### Single-episode predator scent exposure

Rats were exposed to a single episode of predator scent stress in an inescapable environment for 10 min. The day prior to exposure, each rat was assigned to its own exposure cage (i.e. a Plexiglas cage with no bedding) and habituated to the cage environment for 30 min. The cage was left soiled after habituation. On the exposure day, the rat was placed in its corresponding cage for a total of 12 min. After 2 min of adaptation, a cotton pad sprayed with red fox urine was placed at the right end of the cage for another 10 mins (Red fox urine, Wild Life Research Center). The pad was put on a wire mesh inside the cage, right below the lid and out of reach of the rat. The control animal was exposed to the same cotton pad without fox urine. During exposure, the rat was recorded by a video camera. Following predator scent exposure, the rat was returned to his home cage and left undisturbed for 6 days.

### Elevated plus maze (EPM)

The EPM test was performed to measure the anxiety level 6 days after trauma exposure. The maze consisted of four perpendicular arms forming a cross and elevated 50 cm from the floor. Two arms were closed, opposed to each other and enclosed by 40-cm high walls. The remaining two arms, also opposed to each other, were completely open. A camera was placed directly above the maze to record the rat movement.

To perform the test, the rat was initially placed in the center of the maze facing one open arm and allowed to freely explore the maze for 5 min. The times spent in the open and closed arms were determined. The ratio of time in open arms versus that in both open and closed arms [i.e. open/(open+closed)] was calculated to provide a measure of anxiety level. A lower score of this ratio indicates a higher level of anxiety. The maze was cleaned with a 70% alcohol solution in water between tests.

### CORT Measurement

Rats (n = 39) had blood samples collected to quantify the hormonal response to stress. Corticosterone (CORT) level was measured immediately before predator scent exposure (i.e. 0 min) and 30 min, 60 min, and 120 min after exposure to quantify the dynamic response to predator scent stress exposure. Initial blood samples were collected through tail nick, with subsequent time point samples collected by disturbing the tail scab. Samples were collected using heparin coated microtubes with capillary tube inserts (RAM Scientific Safe-T-Fill, Nashville, TN, USA). Samples were centrifuged to obtain plasma and stored at -20 °C. The plasma CORT level was then quantified using CORT enzyme-linked immunosorbent assay (ELISA) kit (Enzo life sciences Inc. Ann Arbor, MI. Cat ADI-901-097) according to the manufacturer’s manual. The total CORT response for each animal was quantified by the area-under-the-curve of the CORT response curve, calculated by the summation of the CORT amplitude across all time points with the CORT level at 0 min (i.e. baseline) being subtracted at each time point. This measurement took into account both the amplitude and duration of the CORT response and represented the total output of the hypothalamus-pituitary-adrenal axis.

### Acclimation and rsfMRI data acquisition

All rats were scanned before the trauma exposure experiment. Before imaging, animals were acclimated to a ‘mock’ imaging environment for 7 days. This acclimation protocol was employed to diminish the stress and motion of the subject during scans ^69^. The rat was first briefly anesthetized with 2-4% isoflurane in oxygen (3-5 min). The head was restrained with a bite bar, nose bar and side pads that restricted the head motion. A pair of shoulder bars and a body tube maintained the body in place without impeding ventilation. Animals were allowed to regain consciousness by discontinuing isoflurane after being set up in the restrainer, typically in 10 mins. The rat was then placed in a dark ventilated box and the sound of different MRI sequences was played, presenting an environment mimicking MRI scanning. All rats were given 7 daily acclimation sessions with a gradually increasing acclimation period (15, 30, 45, 60, 60, 60 and 60 min from day 1 to day 7). A similar approach has also been applied by other research groups ^70-72^.

At least 2 days after acclimation, MRI scans were performed in a 7-Tesla scanner interfaced with the Bruker console system (Bruker, Germany). The rat was first briefly anesthetized with 2-4% isoflurane in oxygen and set up in the same restrainer with a built-in birdcage 1H radiofrequency volume coil. Anesthesia was then discontinued, and the rat was allowed to regain consciousness and placed into the MRI magnet bore. rsfMRI data were obtained using a single shot gradient-echo echo-planar imaging sequence with the parameters of repetition time (TR) = 1 sec, echo time (TE) = 13.8 ms, 64 × 64 matrix size, 20 slices with 1 mm thickness, field of view (FOV) = 32 × 32 mm^2^, in-plane resolution = 500 × 500 μm^2^. Each rat was scanned for 2400 EPI volumes per scan session. Anatomical images were also acquired using a fast spin-echo sequence with the parameters of TR = 1500 ms; TE = 8 ms; matrix size = 256 × 256; FOV = 3.2 × 3.2 cm^2^; slice number = 20; slice thickness = 1 mm; RARE factor = 8.

### rsfMRI data processing

rsfMRI images were preprocessed by performing alignment using Medical Image Visualization and Analysis (MIVA, http://ccni.wpi.edu/), motion correction using SPM 12 (http://www.fil.ion.ucl.ac.uk/spm/), spatial smoothing (Gaussian kernel: FWHM = 0.75 mm) and temporal smoothing (band-pass filter, cutoff frequencies: 0.01-0.1Hz), and regression of motion parameters and signals from the white matter and ventricles as described in past studies ^20,22,23,73^. Before performing these steps, EPI volumes with relative framewise displacement (FD) > 0.2 mm and their immediate temporal neighbors were first removed ^74^. In addition, the first 10 volumes of each rsfMRI run were also removed to ensure the magnetization was at steady state.

Functional connectivity analysis was conducted using region-of-interest (ROI) based correlational analysis. This method evaluated the RSFC by quantifying the Pearson correlation coefficient of regionally averaged spontaneous BOLD signals between different brain regions. Seed maps were generated by correlating the regionally averaged seed time course with the time course of each brain voxel. A linear mixed effect model was used to determine the overall effect on RSFC of each group of rats, while removing any random effect from different batches of animals.

## Acknowledgments

The present study was partially supported by National Institute of Neurological Disorders and Stroke Grant R01NS085200 (PI: Nanyin Zhang, PhD) and National Institute of Mental Health Grant R01MH098003 and RF1MH114224 (PI: Nanyin Zhang, PhD) and R37MH058883 (PI: Gregory Quirk).

