## Supplemental Information for "Individual variability in behavior and functional networks predicts vulnerability using a predator scent model of PTSD"

### Supplemental Figures

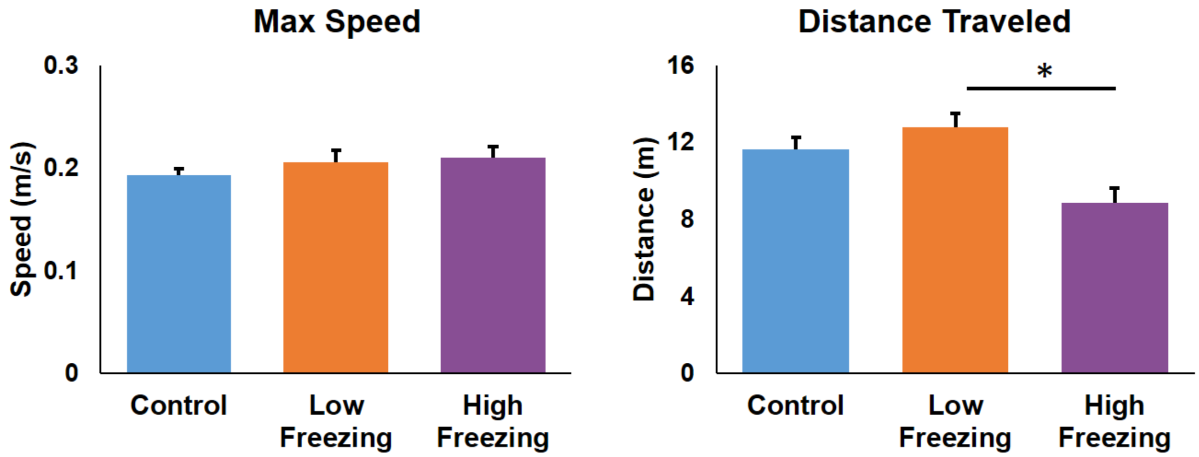


**Supplement Figure 1**. **a**) Maximal speed in high-freezing, low-freezing and control animals. No difference was observed in the three groups (One-way ANOVA, $p = 0.43$). **b**) Distance traveled in high-, low-freezing and control animals. Lower freezers displayed more distance travelled during exposure (One-way ANOVA, $F_{(2,62)} = 8.43$, $p = 0.001$, high freezers vs low-freezers, $p = 0.001$).

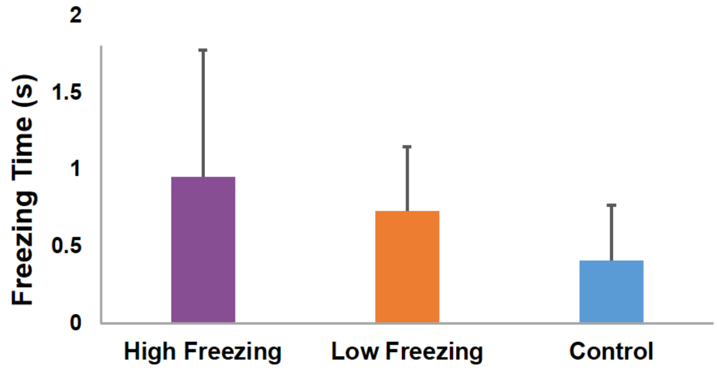


**Supplement Figure 2**. Freezing time during the period of acclimation to the exposure cage immediately before exposure. No group difference was observed (One-way ANOVA, $p = 0.86$).

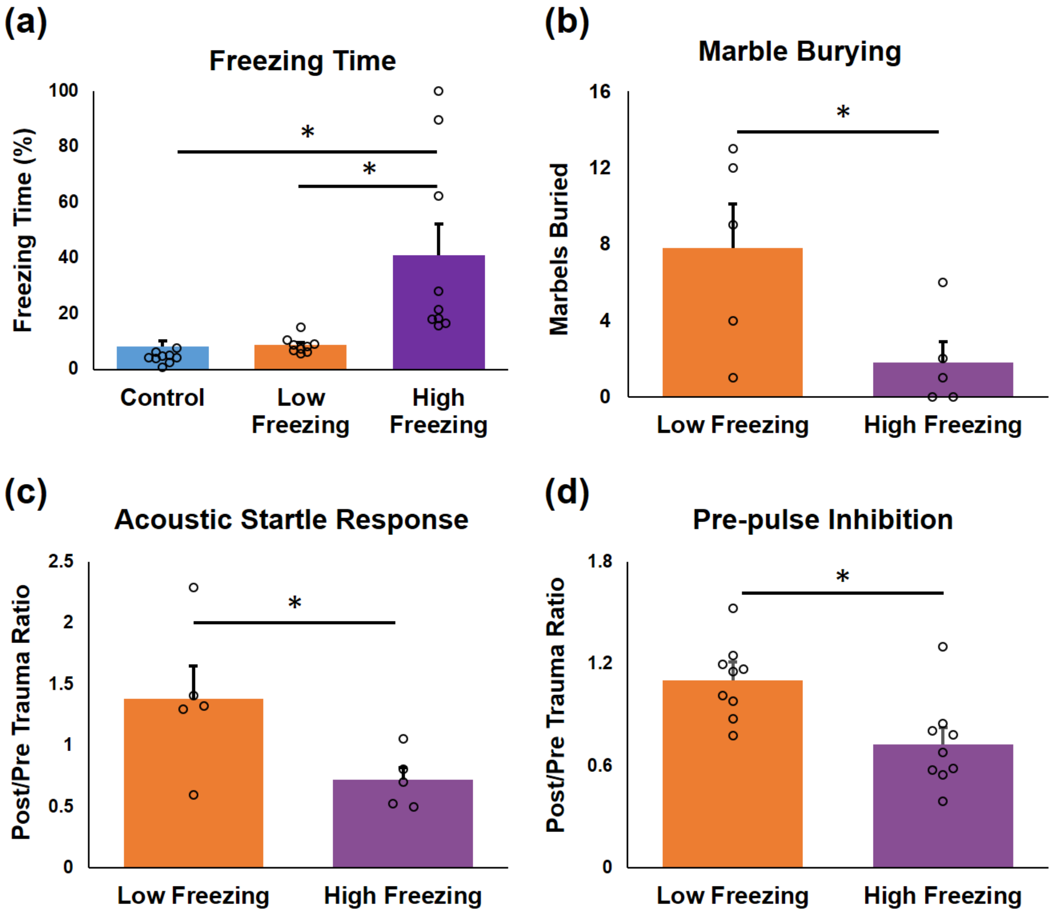


**Supplement Figure 3**. Association with other types of PTSD-like behaviors in low-freezing animals (n = 32, 18 stressed, 14 control). All fox-urine exposed animals were divided into lower-half and higher-half freezing groups (tertiles were not used given the relatively small sample size). **a)** Freezing time in low-freezing, high-freezing and control animals. High-freezing rats showed more freezing (One-way ANOVA, $F_{(2,16)} = 8.24$, $p = 0.011$) than both control ($p < 10^{-9}$) and low-freezing ($p < 10^{-9}$) rats. Consistent with the results in the main cohort of animals, there was no difference in freezing time between control and low-freezing rats. **b)** Low-freezing rats buried more marbles in a marble burying test than high-freezing rats (n = 5 each group, two sample t-test, t = 5.47, p = 0.048) **c)** Acoustic startle response (ASR) in high- and low-freezers. The habituation of ASR was quantified by the averaged response of the last 10 trials normalized by that of the previous 10 trials. The ratio of this quantify 6 days post trauma versus pre trauma was calculated to measure the change in habituation to ASR induced by traumatic stress. This measure was significantly higher in low-freezing animals than high-freezing animals (n = 9 each group, two-sample t-test, t = 5.3, p = 0.050), suggesting that the ability to habituate to acoustic startle stimuli was decreased after trauma exposure in low freezing animals. **d)** Prepulse inhibition (PPI), which measured the suppression of the startle response with a 90 ms prepulse, in low- and high-freezing animals. The ratio of post-trauma PPI versus pre-trauma PPI was significantly greater in low-freezing animals than in high freezing animals (n = 9 each group, two-sample t-test, t = 5.16, p = 0.037), indicating differential impact on sensory gating function by trauma exposure in low-freezing animals.

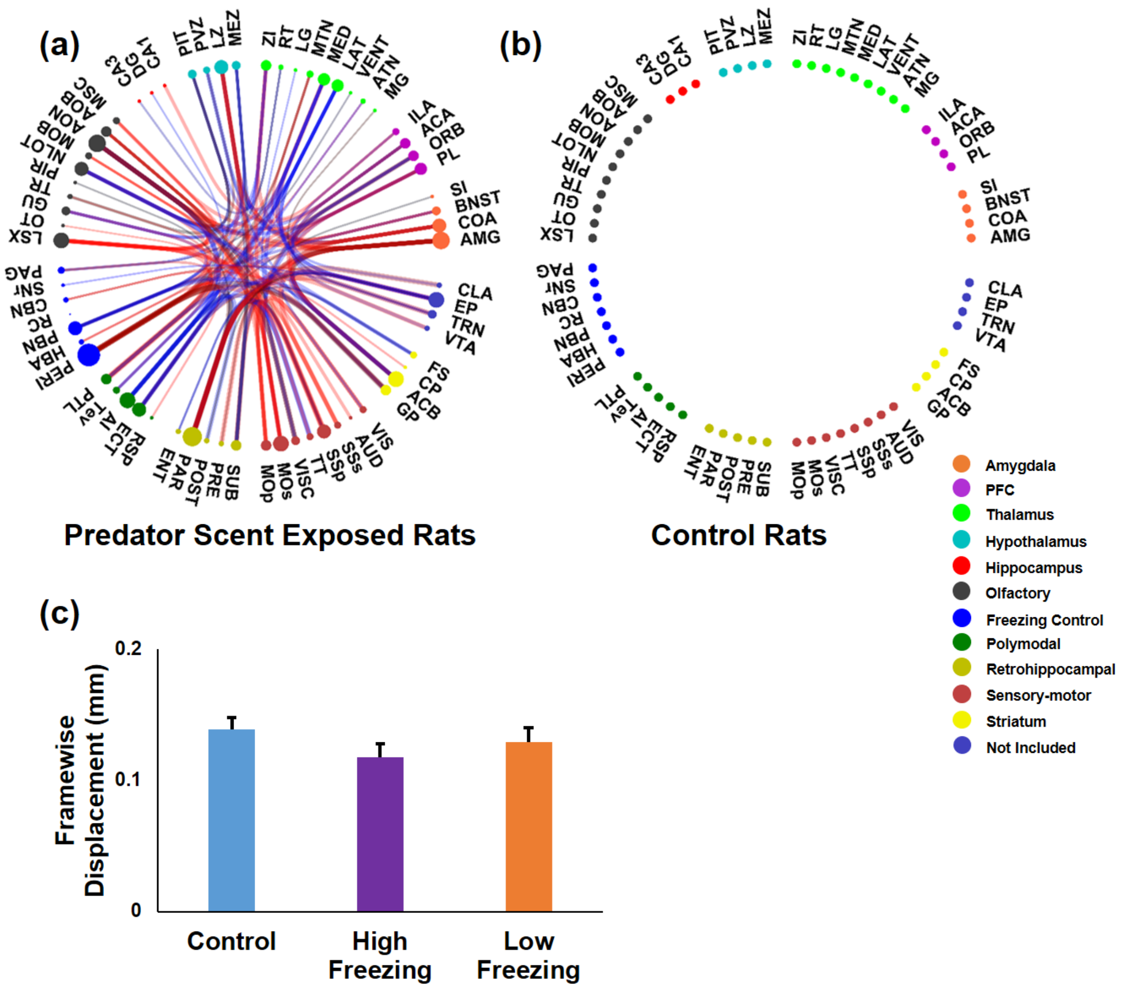


**Supplement Figure 4**. **a**) Neural circuits showing significant difference in pre-trauma resting-state functional connectivity (RSFC) between low-freezing and high-freezing exposed animals. Each line between these circles represents a connection that is significantly different between high-freezing and low-freezing animals. The color represents whether low-freezing animals show increased (red) or decreased (blue) RSFC. The size of each circle around the perimeter is proportional to the number of significant connections displaying significant difference for that region. The transparency of the line is proportional to the strength of that difference. **b**) No circuit displayed pre-exposure RSFC difference between high-freezing and low-freezing control rats. **c**) Motion levels during imaging in low-, high-freezing and control animals, quantified by averaged framewise displacement. No statistical difference in motion levels among three groups was observed ($F_{(2,62)} = 1.23$, $p = 0.30$).

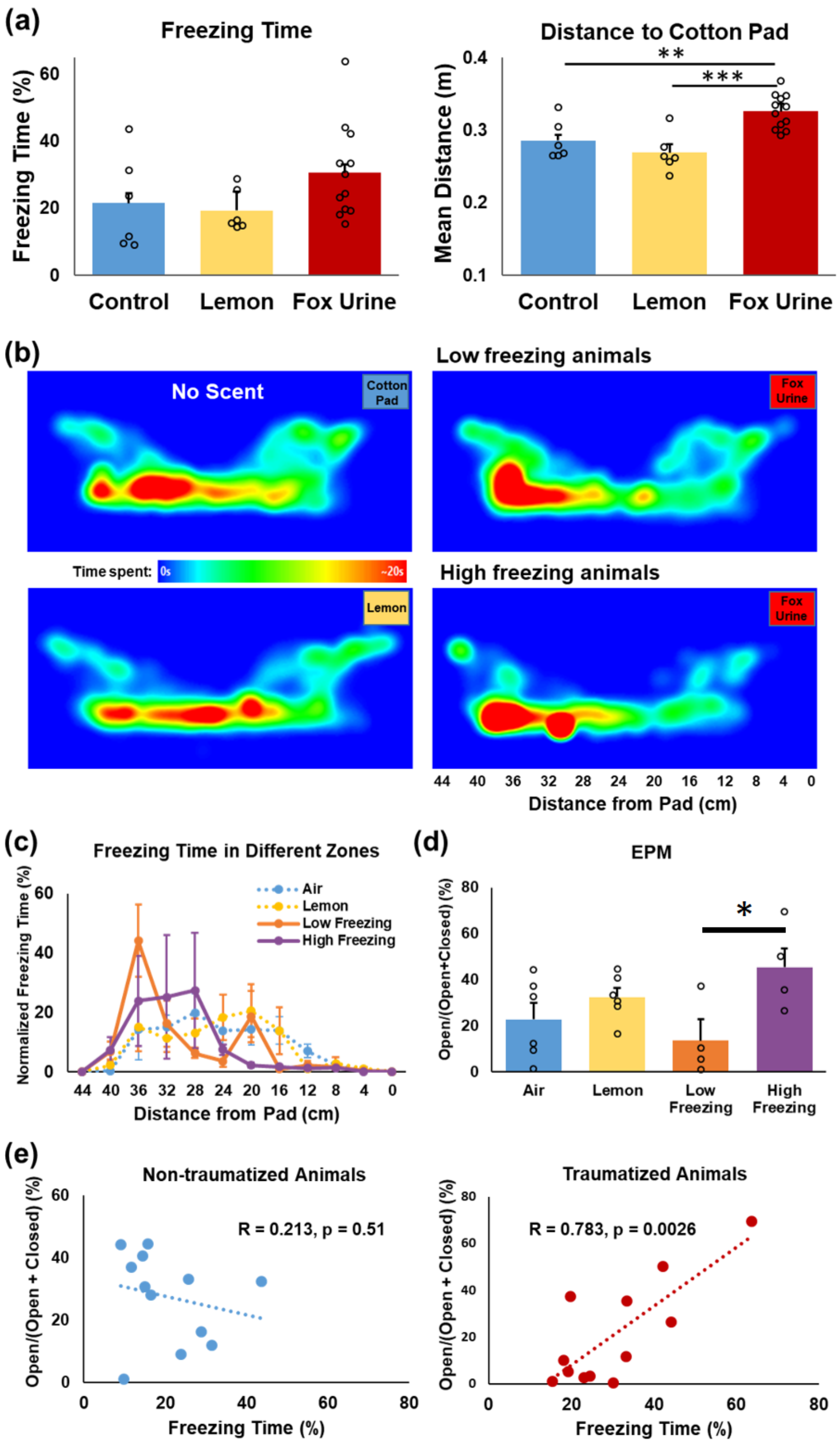

(a)
Freezing Time
Freezing Time (%)
Control
Lemon
Fox Urine
Distance to Cotton Pad
Mean Distance (m)
**
***
(b)
No Scent
Cotton Pad
Low freezing animals
Fox Urine
Time spent:
0s
~20s
Lemon
High freezing animals
Distance from Pad (cm)
(c)
Freezing Time in Different Zones
Normalized Freezing Time (%)
Air
Lemon
Low Freezing
High Freezing
Distance from Pad (cm)
(d)
EPM
Open/(Open+Closed) (%)
*
Air
Lemon
Low Freezing
High Freezing
(e)
Non-traumatized Animals
Open/(Open + Closed) (%)
R = 0.213, p = 0.51
Freezing Time (%)
Traumatized Animals
R = 0.783, p = 0.0026
Freezing Time (%)

**Supplement Figure 5**. **Behavioral response to a control scent (lemon)**. **a**) Lemon exposed animals (n = 6) showed virtually identical freezing (left, t = 2.0, p = 0.16) and avoidance (i.e. distance from the cotton pad, right, t = 1.14, p = 0.32) compared to animals exposed to no scent (i.e. air, n = 6). In contrast, the fox urine exposed group (n = 12) maintained a significantly greater mean distance from the cotton pad (One-way ANOVA, $F_{(2,23)} = 11.61$, $p = 0.0004$) than both air ($p = 0.012$) and lemon ($p = 0.0006$) exposed animals. In addition, fox urine exposed animals showed a trend of more freezing than non-traumatized animals (left, $p < 0.056$, lemon and no-scent groups combined). **b**) Heat maps of the spatial distribution of time spent in the cage for the air-exposed, lemon-exposed, fox urine-exposed low freezing, and fox urine-exposed high freezing groups, respectively. The no-scent and lemon groups showed no evident bias for either side of the cage, while the low-freezing group showed a strong bias toward the far end (from the pad) of the cage. High freezing rats also showed a bias toward the far end of the cage, but to a lesser extent. **c**) The spatial distribution of the normalized freezing time for each group in the cage. Similar to the distributions of the total time spent in different zones, the normalized freezing time in no-scent and lemon rats displayed little bias for either side of the cage. High and low freezing animals showed a spatial bias of normalized freezing time toward the far end of the cage, with an even stronger bias in low-freezing animals. **d**) EPM performed 6 days post exposure in all four groups of animals. Low freezing animals showed a significantly lower EPM score (One-way ANOVA, $F_{(2,17)} = 4.2$, $p = 0.033$) compared to high freezing animals ($p = 0.026$). **e**) The freezing time was positively correlated to the EPM score across all fox urine exposed animals ($r = 0.783$, $p = 0.0026$), but not in non-traumatized animals ($r = 0.213$, $p = 0.506$).

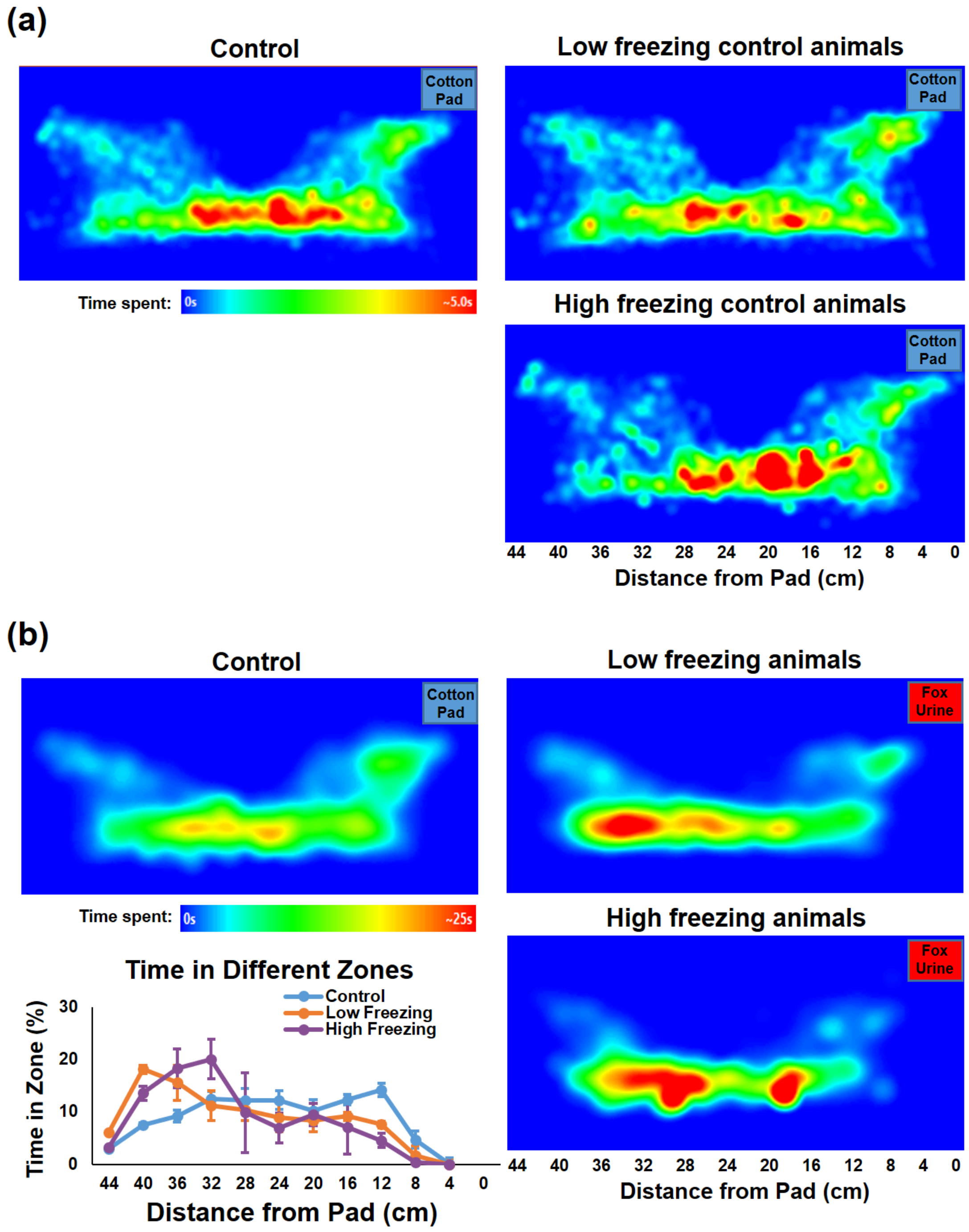


**Supplement Figure 6. Potential impact of acclimation/imaging on fear responses. a**) Heat maps of time spent at each location for all control, low-freezing control and high-freezing control animals. All control animals were divided into tertiles based on their cumulative freezing time. No difference in the distribution of time spent in the cage between low-freezing control and high-freezing control animals was observed (two-way ANOVA, $F_{2,28} = 2.20$, $p = 0.13$). **b**) Heat maps and total time spent at different locations in animals that were not acclimated or imaged (control, n = 20, fox urine exposed, n = 35). Similar differential response during exposure was found in these animals, with lower freezers generally exhibiting more avoidance from the fox urine.
